# Socioeconomic Status and Subdivisions of the Amygdala and Hippocampus in Children and Adolescents

**DOI:** 10.1101/2023.03.10.532071

**Authors:** Jamie L Hanson, Dorthea J Adkins, Brendon M Nacewicz, Kelly R Barry

## Abstract

Socioeconomic status (SES) in childhood can impact behavioral and brain development. Past work has consistently focused on the amygdala and hippocampus, two brain areas critical for emotion, memory, and learning. While there are SES differences in total amygdala and hippocampal volumes, there are many unanswered questions in this domain connected to neurobiological specificity and whether these effects vary by participant age or sex. To address these gaps, we combined multiple large neuroimaging datasets of children and adolescents with information about neurobiology and SES (N=2,765). We examined subdivisions of the amygdala and hippocampus, derived from Freesurfer, using linear mixed effects models. Higher SES was associated with larger volumes across all three amygdala subdivisions examined, the superficial cortical division, basolateral complex, and centromedial region. Within the hippocampus, SES was specifically related to volumes in the head, with no significant associations for the body or tail. Contrary to our hypotheses, we found no significant interactions between SES and participant age or sex after correcting for multiple comparisons, suggesting these associations were relatively consistent across the developmental period examined (ages 5-18) and similar for males and females. These results fill in important gaps regarding the neurobiological specificity of SES effects, demonstrating associations across functionally distinct subdivisions of these critical brain structures.

## INTRODUCTION

Socioeconomic status (SES) in childhood has been associated with multiple negative physical and mental health outcomes, with several meta-analyses noting these links (Peverill et al., 2021; Piotrowska et al., 2015). These disparities, in part, reflect structural inequalities in how resources, opportunities, and risks are distributed across socioeconomic strata, including unequal access to quality education, healthcare, safe housing, and economic security (Adler & Newman, 2002; Braveman & Gottlieb, 2014). The mechanisms underlying these relations, however, are poorly understood. An emerging approach leverages precise quantitation of neurobiology to understand SES-gradients of health (Hair et al., 2015, 2022; Hanson & Hackman, 2012). Neuroscientific investigations may allow a more elemental focus, as the brain determines behavioral and physiological responses (McEwen, 2007). This focus may be particularly valuable given the protracted nature of brain development and that the brain is shaped by experiences early in life (Noble & Giebler, 2020).

A growing body of research has found neurobiological alterations in samples exposed to poverty or lower SES conditions (Johnson et al., 2016; Noble & Giebler, 2020). Notably, childhood poverty has been implicated in structural differences across multiple brain regions, with differences in hippocampal and amygdala structure being commonly reported. The link between childhood poverty and smaller hippocampal volumes has been replicated by at least eight research groups (Ellwood-Lowe et al., 2018; Hanson et al., 2011; Luby et al., 2013; McDermott et al., 2019; Noble et al., 2012, 2015; Raffington et al., 2019; Weissman et al., 2023; Yu et al., 2018). However, studies examining the directional impact of childhood poverty on amygdala structure have produced a landscape of heterogeneous results, with reports of larger and smaller amygdalae (Hanson & Nacewicz, 2021). Given these areas’ connections to important socioemotional functions and learning, understanding how poverty may shape these regions could shed light onto the mechanisms of SES-related disparities. The amygdala is a central neural hub for vigilance and processing negative emotions (Adolphs, 2010; LeDoux, 2007). The hippocampus plays a critical role in memory representations and using previously acquired information in service of goal-directed behavior (Murty et al., 2016; Shohamy & Turk-Browne, 2013). As such, these brain areas are critical for emotion and behavioral responding.

While there are SES differences in amygdala and hippocampal volumes, there are many unanswered questions in this domain connected to neurobiological specificity, and for whom these effects may be more pronounced. First, related to neurobiology, while research often treats the amygdala and hippocampus as unitary structures, they are complex and heterogeneous. Different amygdala nuclei have unique connectivity profiles, patterns of developmental changes, and behavioral correlates (Klein-Flügge et al., 2022; LeDoux, 2007; Swanson & Petrovich, 1998). Similarly, the hippocampus is composed of functionally distinct subregions, with differential connectivity and cytoarchitectonics (Gruber & Ranganath, 2019; Strange et al., 2014; Vogel et al., 2020). The posterior hippocampus has been linked more to cognitive functions, while more anterior regions relate to stress and affect (Fanselow & Dong, 2010; Poppenk & Moscovitch, 2011). Second, there may be potential sociodemographic subgroups where the effects are more pronounced; specifically, sex and age may both moderate the impact of SES on neurobiology. Motivated in part by sex disparities in many neuropsychiatric disorders (Hartung & Lefler, 2019), there has been a growing emphasis on sex as a biological variable. After stress exposure, sex differences in neuronal firing, dendritic spines, neurogenesis, and fMRI responsivity have been found in both the amygdala and the hippocampus (Stevens & Hamann, 2012; van Eijk et al., 2020; Yagi & Galea, 2019). These sex differences may be due to different neuroendocrine processes, responses from the environment, or sex chromosome-specific neuroprogramming (Bale & Epperson, 2017; Shansky & Woolley, 2016). Sex may also be related to differential responses to stress exposure, like those associated with lower SES (Bath, 2020). Related to age, there are non-linear trajectories for brain development, with the amygdala and hippocampus increasing in volume in childhood and adolescence, often in sex-specific way (Fish et al., 2020; Herting et al., 2018; Lenroot & Giedd, 2006; Wierenga et al., 2014). The impacts of stress on neurobiology may vary with age and development (Hanson & Nacewicz, 2021). Stress may increase volumes in certain regions early in development, but then relate to “excitotoxic burnout” and smaller volumes later in time. As such, it will be critical to explore connections between neurobiology, age, sex, and SES.

Connected to neurobiological specificity, there is a growing body of past work examining amygdala and hippocampal subdivisions after childhood adversity. For the hippocampus, past projects have commonly reported smaller volumes in Cornu Ammonis (CA) 1 for those exposed to high levels of adversity (Lee et al., 2018; Margolis et al., 2022; Teicher et al., 2012; Yuan et al., 2020); however, results are not perfectly uniform with other studies only finding differences in CA3, and not CA1 (Brody et al., 2017; Malhi et al., 2019). With the amygdala, less work has been completed. Two studies reported smaller basolateral amygdala volumes after exposure to adversity (Nogovitsyn et al., 2022; Oshri et al., 2019), but these projects also reported stress was sometimes related to differences in accessory basal, central-medial, and paralaminar subdivisions. Related to SES, limited work has examined if there are potential alterations in volumes of amygdala and hippocampal subdivisions. Three past studies have found anterior hippocampal volumes (i.e., Dentate gyrus; CA1) were positively related to different operationalizations of SES, including parental education (Merz et al., 2019), family household income (Decker et al., 2020), and socioeconomic conditions in a census tract (Botdorf et al., 2022). Notably, no work to date has examined amygdala subdivisions in relation to SES.

Attempting to overcome these limitations, here, we combined multiple, large neuroimaging datasets of children and adolescents with information about neurobiology and SES (N=2,765). To improve neurobiological specificity, we examined subdivisions of the amygdala and hippocampus. We aimed to richly probe the main effects of SES on these smaller regions, as well as examine potential sex- or age-specific impacts of SES on these volumes. In keeping with past reports of smaller volumes in lower SES youth, we predicted lower SES would be related to smaller volumes in amygdala and hippocampal subdivisions. Related to past human and non-human research in stress-exposed groups (Hanson & Nacewicz, 2021), we also predicted that lower SES would be related to smaller volumes in the head of the hippocampus, as well as the basolateral and central amygdala. Finally, given potential neuroprotective effects of estrogen (McEwen, 2002, 2020) and developmental trajectories of brain development (Aoki et al., 2017), we predicted relations between SES and volumes would be stronger for older participants, especially boys.

## METHOD

### Participants

Participants between 5-17 years of age were drawn from four large neuroimaging projects: The National Consortium on Alcohol and Neuro-Development in Adolescence (NCANDA; (Brown et al., 2015), the Healthy Brain Network (HBN; (Alexander et al., 2017), the Pediatric Imaging, Neurocognition, and Genetics (PING study) (Jernigan et al., 2016), and the Human Connectome Project in Development (HCP-D; (Somerville et al., 2018). Richer sample descriptions are available in our supplemental materials. Across these studies, the total number of participants with usable data was N=2765 (44% Female; Mean Age= 11.9, Age SD= 3.5; Age Range = 5.04-17.99). Descriptive information for the combined sample is shown below in Table 1. A histogram of participant age by study is shown in Figure 1.

**Figure 1.**
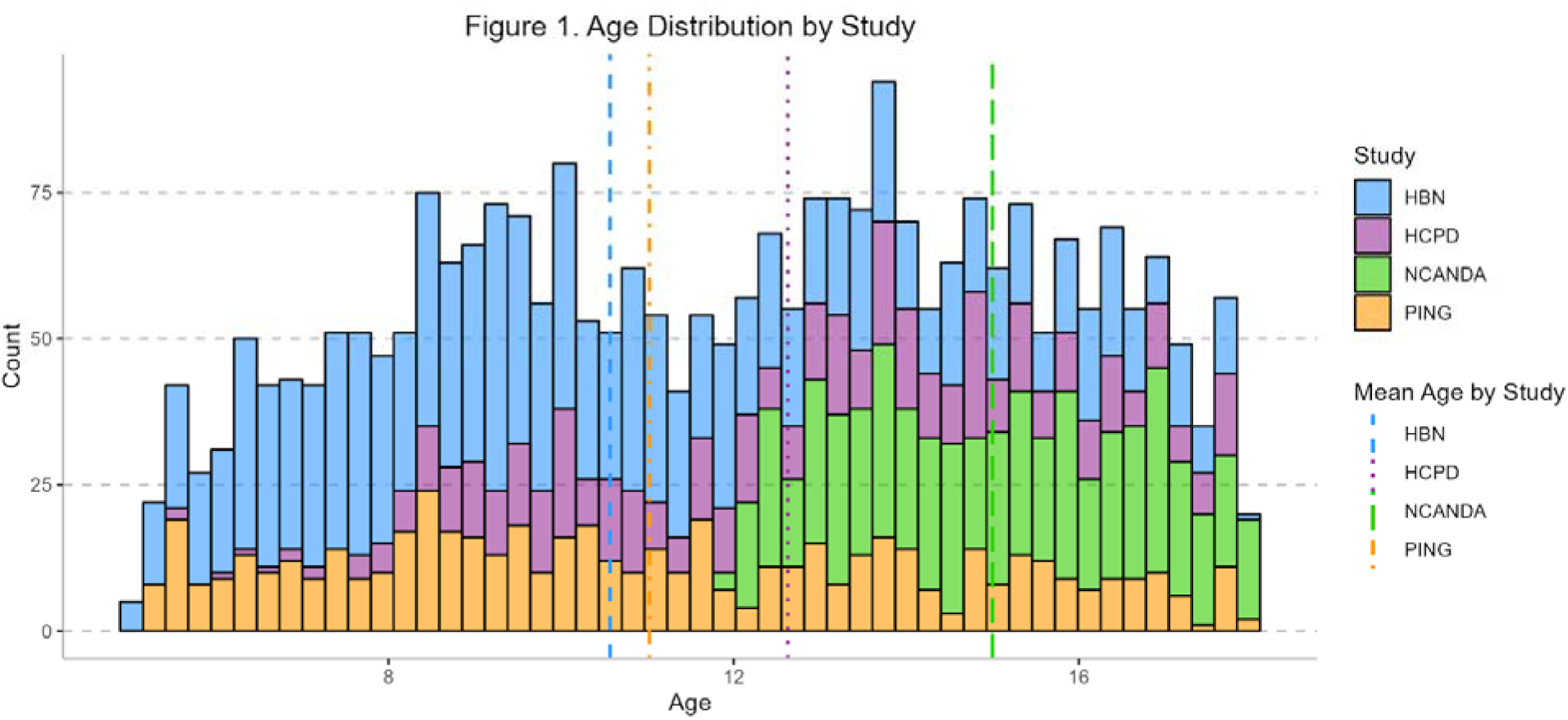
Stacked histogram depicting the number of participants at each age, for each study, between the ages of 5 and 18. The top set of bins represents the numbers of participants in the HBN study at each age (colored in blue); stacked below are the bins for the HCPD (purple), NCANDA (green), and HBN (orange) studies, respectively. The dashed and dotted lines display the mean ages for each study (with each color corresponding to a different study; blue = HBN; purple = HCPD; green = NCANDA; orange = PING).

**Figure 2.**
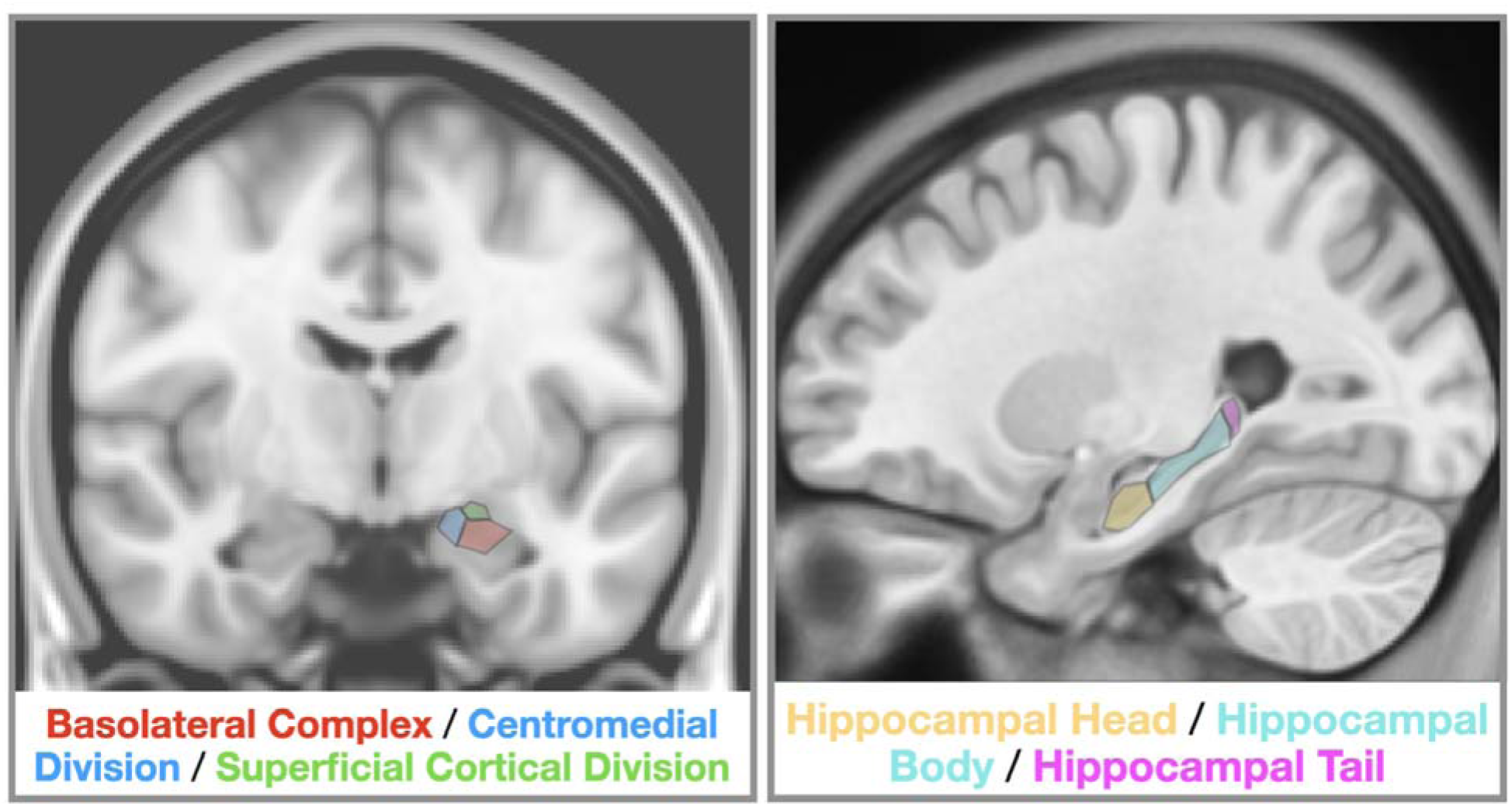
Depiction of our amygdala and hippocampal regions of interest. The left side of this figure shows our 3 amygdala subnuclei groups overlaid on an average T1-weighted scan. The right side of this figure shows our 3 hippocampal subdivisions also overlaid on an average T1-weighted scan.

**Figure 3.**
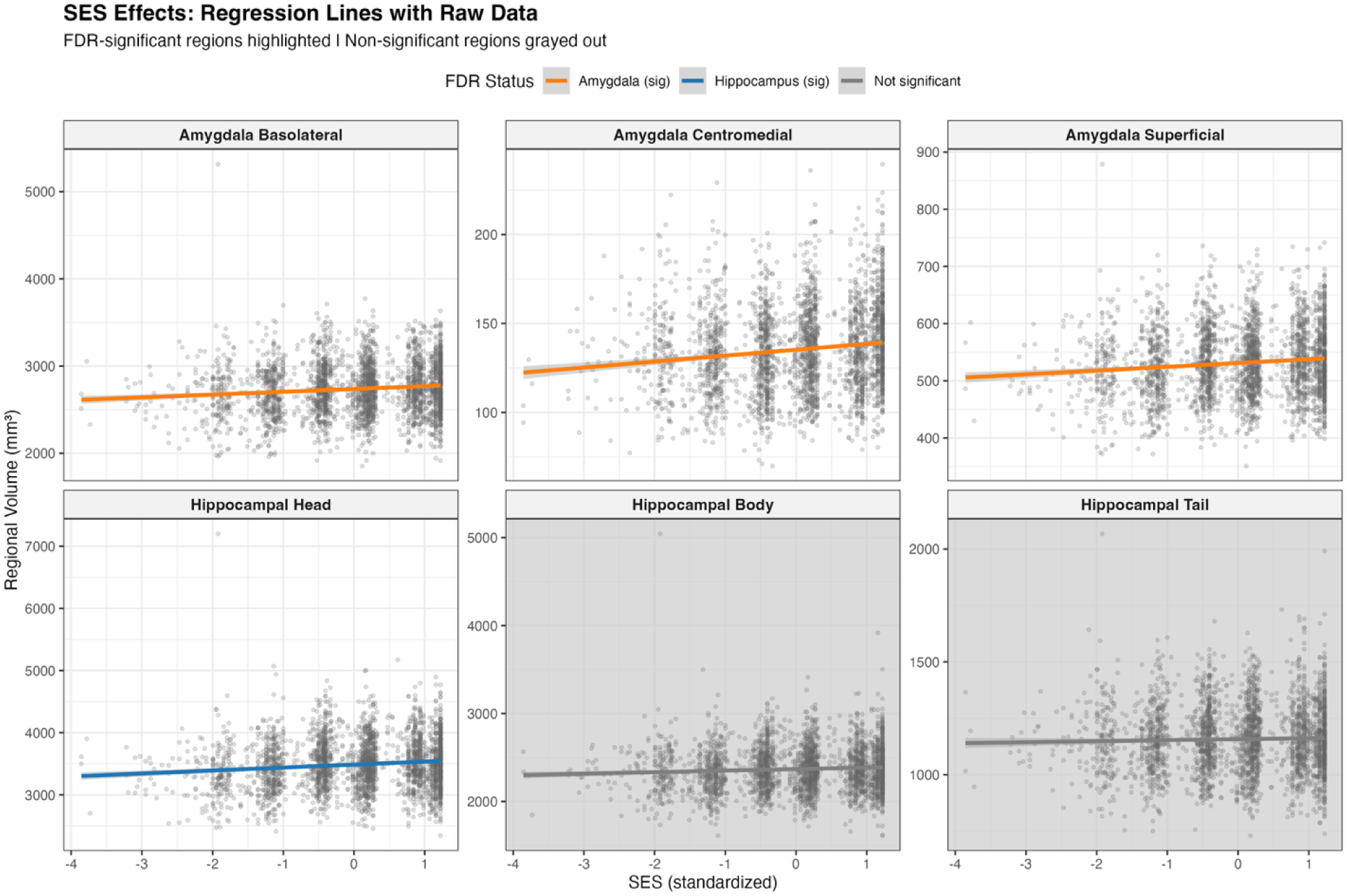
Scatterplots depicting main effects of socioeconomic status on amygdala and hippocampal subdivisions. Each panel displays the relationship between standardized SES composite scores (x-axis) and regional brain volumes in mm³ (y-axis) for six subcortical subdivisions: amygdala basolateral complex, centromedial division, and superficial cortical division (top row), and hippocampal head, body, and tail (bottom row). Orange and blue regression lines (amygdala = orange; blue = hippocampus) indicate FDR-significant positive associations between SES and regional volumes (q < 0.05). Gray lines indicate non-significant associations. All models controlled for age, sex, estimated total intracranial volume, image quality, and study site.

**Table 1.** Sample demographics by study. Demographic information for the full sample and by study, including means and standard deviations for age (in years), highest caregiver education (in years), image quality (CAT12 grade), and estimated total intracranial volume (mm³); median and interquartile range for family income (USD); and the distribution of sex across studies.

| Variable | Overall<br>(n = 2765) | HBN<br>(n = 1184) | HCPD<br>(n = 460) | NCANDA<br>(n = 566) | PING<br>(n = 555) | p-value |
| --- | --- | --- | --- | --- | --- | --- |
| **Age (years), Mean (SD)** | 11.91 (3.45) | 10.57 (3.28) | 12.63 (2.87) | 15.00 (1.69) | 11.02 (3.46) | <0.001 |
| **Sex, n (%)** |  |  |  |  |  | <0.001 |
| Male | 1,537 (56%) | 748 (63%) | 209 (45%) | 282 (50%) | 298 (54%) |  |
| Female | 1,228 (44%) | 436 (37%) | 251 (55%) | 284 (50%) | 257 (46%) |  |
| **Family Income, Median [IQR]** | 124,999.50<br>[62,499.50,<br>195,000.00] | 124,999.50<br>[64,999.50,<br>200,000.00] | 116,500.00<br>[75,000.00,<br>175,000.00] | 149,999.50<br>[62,499.50,<br>149,999.50] | 74,999.50<br>[44,999.50,<br>124,999.50] | <0.001 |
| **Highest Caregiver Education, Mean (SD)** | 17.49 (2.95) | 18.67 (2.99) | 16.51 (2.18) | 16.84 (2.47) | 16.45 (2.99) | <0.001 |
| **Image Quality (CAT12 Grade), Mean (SD)** | 0.86 (0.02) | 0.87 (0.02) | 0.84 (0.01) | 0.84 (0.01) | 0.85 (0.01) | <0.001 |
| **Estimated Total Intracranial Volume (mm<sup>3</sup>), Mean (SD)** | 1,501,276<br>(172,642) | 1,492,523<br>(162,581) | 1,595,849<br>(147,332) | 1,486,421<br>(172,252) | 1,456,712<br>(184,962) | <0.001 |

Note. HBN = Healthy Brain Network; HCPD = Human Connectome Project Development; NCANDA = National Consortium on Alcohol and Neurodevelopment in Adolescence; PING = Pediatric Imaging, Neurocognition and Genetics. p-values from Kruskal-Wallis tests (continuous variables) and Fisher's exact test (sex).

### MRI Data Acquisition and Imaging Processing

High-resolution T1-weighted structural images were acquired with varying parameters across each project. The majority of scans came from 3T scanners, with the exception of a 1.5T scanner used at one site. Scans resolution varied in in-plane resolution from 0.8 to 1.2mm. Information about MRI parameters are noted in our supplemental materials. These MRI scans were processed in Freesurfer 7.1, deployed via Brainlife.io (Hayashi et al., 2024). Freesurfer is a widely documented morphometric processing tool suite, http://surfer.nmr.mgh.harvard.edu/ (Dale et al., 1999; Fischl et al., 2002). Based on hand-tracing on high-definition, ex-vivo T1-weighted 7T scans, Freesurfer can output 12 subfields and 9 amygdala subnuclei (see (Iglesias et al., 2015; Saygin et al., 2017) for additional details). With the hippocampus, we focused on segmentation that divided this region into the head, body, and tail. This was motivated by: 1) work finding that hippocampal organization and connectivity varies on its longitudinal axis (i.e., head/body/tail) (Genon et al., 2021; Vogel et al., 2020); and 2) commentary suggesting that segmentation of the longitudinal axis of the hippocampus is more appropriate, than much smaller (and potentially less reliably) segmented subfields, i.e., dentate gyrus, and subiculum (Kahhale et al., 2023; Wisse et al., 2021). With the amygdala, subnuclei volumes were grouped into three divisions based on cytoarchitecture, connectivity, and functional organization (Amunts et al., 2005; Saygin et al., 2017): (1) a basolateral complex (lateral, basal, and accessory basal nuclei), comprising glutamatergic principal neurons that serve as the primary input station for cortical and thalamic sensory information and support associative learning (Paton et al., 2006; Sah et al., 2003); (2) a centromedial division (central and medial nuclei) consists predominantly of GABAergic output neurons that project to hypothalamic and brainstem autonomic centers to coordinate neuroendocrine and behavioral stress responses (Ciocchi et al., 2010); (3) a superficial cortical division (cortical nucleus, anterior amygdaloid area, and corticoamygdaloid transition area) represents transitional cortical-like regions with prominent olfactory and limbic connectivity (Kemppainen et al., 2002; Majak & Pitkänen, 2003). The paralaminar nucleus was excluded from analyses due to ongoing debates about subgroups (deCampo & Fudge, 2012; Kedo et al., 2018), as well as segmentation uncertainty and inconsistent detection across participants (Kahhale et al., 2023). For the 3 hippocampal and the 3 amygdala subdivision groups, we calculated the total volume by summing volumes from the left and right hemispheres of each of these hippocampal or amygdala subdivisions.

To exclude particularly high-motion scans and limit the impact of image quality on subcortical segmentation, we generated a quantitative metric of image quality combining noise-to-contrast ratio, coefficient of joint variation, inhomogeneity-to-contrast ratio, and root-mean-squared voxel resolution (Gaser et al., 2024). Scans with an image quality grade < 0.8 were excluded from analyses, a threshold determined to be effective in previous projects for ensuring adequate volumetric and morphometric measurements (Adkins & Hanson, 2025; Bacas et al., 2023; Gilmore et al., 2021; Hanson et al., 2024). The image quality metric was then included as a covariate in all statistical models to further account for remaining variability in scan quality among retained scans. This was motivated by our work finding that T1-weighted image quality is related to Freesurfer outputs (Gilmore et al., 2021). Additional details about this metric, image processing in Freesurfer, and MRI Data Acquisition are noted in our supplemental materials.

### Measures of SES

We used a multimethod determination of SES, using metrics of both caregiver education and household income. This was motivated by the understanding that income and education each capture distinct facets of socioeconomic position, and that their combination yields a more reliable index of the underlying construct. For caregiver education, caregivers reported on how much school they completed (e.g., obtained a high school diploma; some college; graduate degree); this was converted into numbers of years (i.e., high school diploma = 12 years; some college = 14 years) and then took the highest value by either caregiver. For household income, caregivers selected an income range (i.e., $30,000-39,999/year; $50,000-$99,999/year), or reported a continuous value. In keeping with recommendations from demographers (Parker & Fenwick, 1983), we took the midpoint for reported ranges and then log-transformed this value (i.e., $30,000-39,999=$34,999.5; log($34,999.5)=4.54406184). For continuous income reports, we also log-transformed these values. Then, we created an SES composite by taking an average of z-scored, log-transformed income and z-scored, maximum parental education. In our supplement, we examined income and parental education independently.

### Analytic plan

To test relations between SES and volumes of the hippocampus and amygdala, we employed linear mixed effects models (LMEMs) using the *lme4* R package (Bates et al., 2015). In all models, we included a random effect for study site, and examined the independent (fixed effect) variables of participant age (in years), participant sex (binary-coded), estimated Total Intracranial Volume (eTIV), image quality (CAT12 rating), our SES composite, and interactions between age, sex, and SES. These models include two- and three-way interactions of these variables (i.e., SES x Age x Sex; SES x Sex) and were completed for 6 regions of interest (3 hippocampal subdivisions [head, body, and tail]; 3 amygdala subdivisions [accessory basal, central amygdala, basolateral complex, and medial cortical]). We corrected these different steps of analyses for multiple comparisons using False Discovery Rate approaches (Benjamini & Hochberg, 1995). For any models yielding significant interactions, we planned to probe these effects using simple slopes analysis to examine the association between SES and regional volumes at different levels of the moderating variable (e.g., testing the SES-volume relationship separately for males and females, or at ±1 SD of age). Of note, we did not apply ComBat harmonization, a common approach to dealing with neuroimaging data from different sites. This choice was motivated by recent work finding that harmonization methods obscured some of the true biological effect in structural data and structural MRI data “was harmed by harmonization” (Richter et al., 2022). The mixed-effects modeling approach allowed us to statistically control for systematic site-level differences while maximally preserving individual-level variation in brain structure. In our supplement, we also completed analyses: 1) focused on volumes and household income or education (as individual/separate variables); 2) for smaller parcellations of the hippocampus and amygdala output by Freesurfer; and 3) sensitivity analyses using Generalized Additive Mixed Models and leave-one-site-out cross-validation.

## RESULTS

### Main Effects of SES on Amygdala and Hippocampal Subregions

SES was associated with volumes of all amygdala subdivisions and the hippocampal head. For the amygdala, higher SES was related to larger volumes in the superficial cluster (β*=0.07, p<0.001*), basolateral complex (β*=0.05, p=0.014*), and centromedial region (β*=0.05, p=0.028*). Among hippocampal subfields, SES was only related to volumes in the head of the hippocampus (β*=0.07, p=0.002*), with no significant associations observed for the body or tail (p’s >.7). After correcting for multiple comparisons, all significant associations remained robust (*superficial: p-fdr=0.005; hippocampal head: p-fdr=0.006; basolateral: p-fdr=0.028; centromedial: p-fdr=0.042*).

We examined whether the relationship between SES and regional brain volumes varied as a function of participant age. After correcting for multiple comparisons, no significant SES × Age interactions emerged for any of the amygdala or hippocampal subdivisions (all p-fdr>0.38). The strongest effect was observed in the hippocampal tail (β=0.04, p=0.095), suggesting a trend toward stronger positive SES associations with increasing age, though this did not reach statistical significance. Overall, the association between SES and subcortical volumes did not differ significantly across the age range examined in this sample.

### Two-Way Interaction: SES × Sex

To determine whether sex moderated the relationship between SES and regional brain volumes, we tested for SES × Sex interactions across all subdivisions. No significant interactions were observed for any amygdala or hippocampal subdivision after FDR correction (all p-fdr>0.70). The largest effect was in the amygdala superficial cluster (β=-0.03, p=0.233), though this was far from significant. These null findings indicate that the association between SES and subcortical volumes was similar for males and females in this sample.

### Three-Way Interaction: SES × Sex × Age

Finally, we investigated the three-way interaction between SES, sex, and age to test whether sex-specific SES effects varied across development. No three-way interactions reached statistical significance after FDR correction (all p-fdr>0.19). While the hippocampal tail (β=-0.06, p=0.060) and amygdala centromedial region (β=-0.06, p=0.064) showed marginal trends, these did not survive correction for multiple comparisons. Overall, there was no evidence that the relationship between SES and subcortical volumes varied as a function of both sex and age.

### Main Effects of Age and Sex on Subcortical Volumes

Independent of SES and consistent with prior developmental neuroimaging research, we observed robust main effects of age and sex on subcortical volumes. Age demonstrated significant positive linear associations with volumes across all amygdala and hippocampal subdivisions (*all p-fdr<0.001*), with the strongest effects in the hippocampal body (β=0.12) and tail (β=0.11). Additionally, significant quadratic age effects were observed for all regions (*all p-fdr<0.001*), indicating inverted-U developmental trajectories with volume increases during childhood followed by decreases in later adolescence. Main effects of sex were also pronounced, with females showing larger volumes than males across all three amygdala subdivisions (basolateral: β=0.10, p-fdr<0.001; centromedial: β=0.11, p-fdr<0.001; superficial: β=0.08, p-fdr=0.003) and the hippocampal head (β=0.08, p-fdr=0.002). Sex effects were not significant for the hippocampal body or tail after FDR correction (p-fdr>0.10). Figure 4 displays the pattern of standardized coefficients for all main effects and interactions across the examined subdivisions.

**Figure 4.**
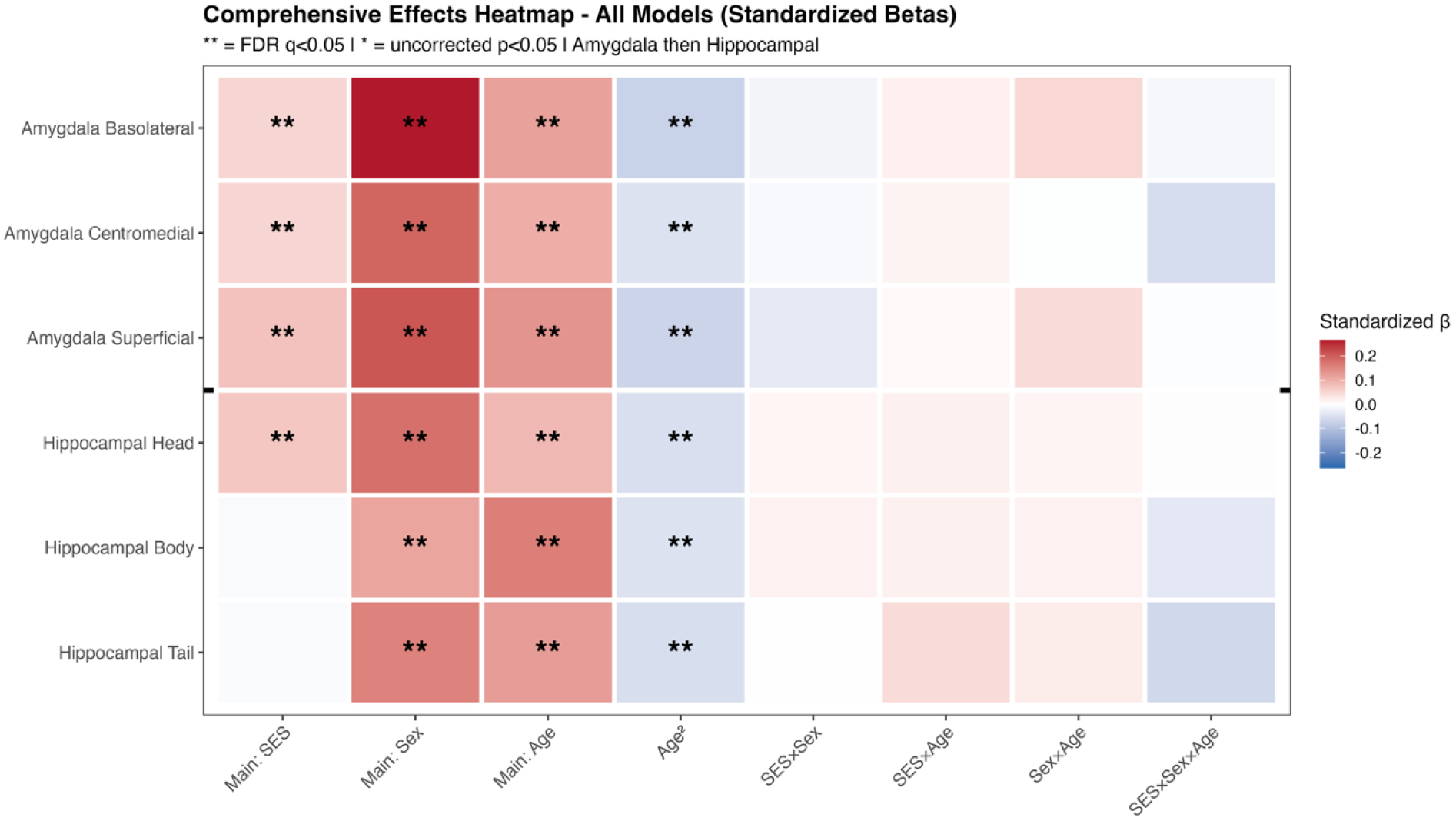
Comprehensive effects heatmap of socioeconomic status, sex, and age associations with subcortical volumes. Heatmap displays standardized regression coefficients (β) for main effects (SES, Sex, Age, Age²), two-way interactions (SES×Sex, SES×Age, Sex×Age), and three-way interaction (SES×Sex×Age) across amygdala subdivisions (top three rows: basolateral complex, centromedial division, superficial cortical division) and hippocampal subdivisions (bottom three rows: head, body, tail). Red shading indicates positive associations; blue shading indicates negative associations; white indicates no association. Double asterisks (**) denote effects surviving False Discovery Rate correction (q < 0.05); single asterisks (*) indicate uncorrected significance (p < 0.05). All models controlled for estimated total intracranial volume (eTIV), image quality (CAT12 rating), and study site (modeled as a random effect).

### Supplemental Analyses

As noted in our supplemental materials, income and education each reproduced the pattern when modeled separately (income more broadly), and the finer FreeSurfer parcellations localized the associations to head-region hippocampal subfields and mostly basal amygdala subnuclei. SES main effects were similar when using GAMMs to model age non-linearly, and leave-one-site-out cross-validation recovered the amygdala and hippocampal-head effects in all 21 folds. No three-way interaction survived multiple comparison correction.

## DISCUSSION

This study investigated associations between SES and volumetric variations in the hippocampus and amygdala. We parcellated the hippocampus and amygdala into smaller subdivisions, aiming to increase neurobiological specificity. With the amygdala, we found that SES was positively associated with volumes across all three subdivisions investigated--the basolateral complex, centromedial region, and superficial cortical division. For the hippocampus, effects were localized to the head of the hippocampus, with higher SES being associated with larger volumes in that subdivision. We tested whether these associations varied by participant age or sex through two-way and three-way interaction models. However, we did not find significant moderation effects after correcting for multiple comparisons. Our findings are partially in line with our a priori predictions. We predicted that subdivisions of the hippocampus and amygdala would be related to SES, but we also believed that relations would vary by sex and age; however, interaction analyses did not support these hypotheses in this large sample.

It is critical to note that these SES-related neurobiological differences likely reflect structural and systemic inequalities rather than inherent biological differences between socioeconomic groups. The volumetric variations we observe are likely downstream consequences of differential access to resources, opportunities, and environmental quality--disparities that are fundamentally rooted in structural inequalities including unequal distribution of wealth, residential segregation by income, educational funding disparities, limited access to quality healthcare, and neighborhood-level disadvantage (Braveman & Gottlieb, 2014; Chetty et al., 2014; Jones et al., 2025; Miller et al., 2025, 2026). These structural barriers constrain opportunities for upward mobility and create conditions where lower-income families face chronic stress, resource scarcity, and reduced access to enriching environments--factors that may shape the neurodevelopmental patterns we observe.

With the hippocampus, our results are similar to previous projects that have found relations between SES and volumetric differences in the whole hippocampus (Hair et al., 2015; Hanson et al., 2011, 2015; Luby et al., 2013; Noble et al., 2012, 2015). With subfields and specific subdivisions, our results are mostly in line with the small number of publications focused on this question. However, each of these papers varies in the parcellation of the hippocampus and the SES variable examined. Specifically, Merz and colleagues found lower parental education was related to smaller volumes in the dentate gyrus and CA1 subfields of the hippocampus (Merz et al., 2019). The CA1 subfield is squarely in the anterior (head) portion of the hippocampus, while their dentate gyrus subfield can span the head and body of the hippocampus. Botdorf and coworkers found smaller anterior and posterior volumes in the hippocampus with greater area deprivation index, a census-based index of SES (Botdorf et al., 2022). Decker et al. found that lower household income was related to smaller anterior hippocampal volumes (Decker et al., 2020). Examined collectively, there is clear evidence that facets of SES are related to volumes in the head of the hippocampus.

Regarding the amygdala, there is a raft of inconsistencies in past work focused on amygdala volumes, SES, and stress exposure (see (Hanson & Nacewicz, 2021). Multiple groups have reported smaller (whole) amygdala volumes in lower SES youth, but results have not been perfectly uniform. There has been no published work to date focused on SES and amygdala subnuclei. In adult samples exposed to childhood adversity, smaller volumes of basal and accessory basal portions of the amygdala have been reported (Nogovitsyn et al., 2022; Oshri et al., 2019). Oshri and colleagues also found high levels of adversity were related to smaller volumes in central-medial portions of the amygdala. Our findings extend these relations to multiple amygdala subdivisions in a large pediatric sample, demonstrating that SES-related volumetric differences are evident across functionally distinct amygdala regions including the basolateral complex (which serves as the primary sensory input station), the centromedial region (which coordinates autonomic stress responses), and the superficial cortical division (which maintains prominent olfactory and limbic connectivity).

Related to age- and sex-specific effects, we tested whether the association between SES and subcortical volumes varied as a function of participant age, sex, or their combination. Despite a reasonably large sample (N=2,765), we found no significant two-way or three-way interactions after FDR correction. This suggests that the relations between SES and volumes were similar for males and females and across the age range. We should, however, interpret these null with care, as interaction analyses are often underpowered even in large samples (Baranger et al., 2023; Castillo et al., 2025; McClelland & Judd, 1993). Past work has provided theoretical rationale for expecting age- and sex-specific effects, but subtle moderation effects could exist below our detection threshold. Regarding sex-specific effects, limited work has specifically considered relations between SES and these volumes in males versus females, with inconsistent findings across studies focused on (Kim et al., 2019; Rakesh et al., 2021). Preclinical work examining “sex as a biological variable” suggests that stress exposure often causes more significant neurobiological alterations in males compared to females (Shansky & Woolley, 2016), with chronic stress causing dendritic atrophy and spine elimination in male rodents (McEwen, 2001), but only slight changes in female rats (Galea et al., 1997). With age, Merz et al. found lower SES was significantly associated with smaller amygdala volumes in adolescent participants, but there were no significant associations for younger age participants (Merz et al., 2018). Previously, we also noted that in adversity exposed samples larger volumes in the amygdala were seen for child samples, but small volumes were reported in adults (Hanson & Nacewicz, 2021). Additionally, some evidence suggests timing matters: adversity exposure before 8 years of age was more likely to impact hippocampal volumes in males, while adversity after 9 years of age impacted females more significantly (Teicher et al., 2018). Interactive effects may exist, but our sample design and analytical approaches may have made detection more challenging. Future work, especially longitudinal studies, will be critical to providing clarity about potential age- and sex-specific neurobiological impacts of SES.

Given our results, it will be crucial to think about how these neurobiological alterations may influence future behavior. We did not explore relations between volume and behavior, as specific behavioral measures greatly varied by project. In thinking about relations between SES and the head of the hippocampus, a growing literature suggests that this hippocampal subdivision may be more related to emotion and stress responding (Herman et al., 1998; Kheirbek et al., 2013; Strange et al., 2014). Lesions to this region in rodents (Henke, 1990), as well as variations in human functional connectivity of this area (Gryglewski et al., 2019; Hanson, Gillmore, et al., 2019; Hubachek et al., 2021) have been linked to alterations in stress responding and emotional behavior. Thinking about the amygdala, we find multiple subdivisions are related to SES and these different divisions have been implicated in different aspects of fear responsivity and reward valuation (Phelps & LeDoux, 2005). The basolateral complex serves in sensory gating and associative learning, while the centromedial division relays emotional information to coordinate autonomic responses. Collectively, these alterations could create attentional biases toward negative valenced stimuli, potentially contributing to challenges in emotion regulation (Barry et al., 2022; Hanson, Albert, et al., 2019; Oshri et al., 2019). Preclinical work supports these broad presumptions, as mice with smaller basolateral volumes show significantly greater levels of conditioned freezing compared to those with larger volumes (Yang et al., 2008).These different behavioral processes will be important to interrogate in future work connecting SES and medial temporal lobe neurobiology.

Considering the strengths of our work, the current study benefited from a large sample size (N=2,765), well-validated methods, and an SES composite considering both parental education and income. However, we must consider potential issues with the work. First, our work was cross-sectional in nature, based on a single MRI scan. Volumetric differences could “equalize” over time; this may be particularly true of the hippocampus, where research has demonstrated reversibility in volumetric differences if given a “stress-free” period (Vyas et al., 2002). In future work, we hope to assess other structural and functional properties of the amygdala and hippocampus through the use of longitudinal functional MRI and magnetic resonance spectroscopy (Nacewicz et al., 2012). Second, our data were pooled across many sites and acquisition protocols. We modeled site as a random effect, but per-scanner identifiers were not consistently available across datasets, so we could not separate scanner from site or account for within-site scanner changes (e.g., a mid-study upgrade). Because acquisition is organized at the site level, the site term should capture most scanner-related variance. Third, automated methods, like FreeSurfer, may be less accurate than hand-delineation of these subdivisions (Hanson et al., 2012). Quantification of these areas by hand is not feasible in such a large sample, but there may be novel automated approaches that more accurately quantify small amygdala and hippocampal subdivisions (DeKraker et al., 2022; Liu et al., 2020). Fourth, our null findings for age and sex interactions should be interpreted with appropriate caution.

Interaction analyses require substantially larger sample sizes than main effect analyses to achieve adequate power (McClelland & Judd, 1993), and subtle moderation effects could exist below our detection threshold despite our large sample. Finally, it will be important to think mechanistically about how lower SES may be impacting neurobiology. Lower SES encompasses a host of challenges likely to impact development, including higher levels of stress, food insecurity, residential instability, community violence, and structural disadvantage (Evans, 2004; Hanson & Hackman, 2012; Masarik & Conger, 2017). This fits with past work finding relations between hippocampal volumes and rich measures of stress exposure (Hanson et al., 2015). Similarly, environmental stimulation is another proximal factor through which SES may impact these volumes, especially the hippocampus. Economically marginalized families may have lower levels of cognitive stimulation in the home, including fewer toys and educational resources (Evans, 2004; Masarik & Conger, 2017). Environmental enrichment and stimulation can impact hippocampal structure, including dendritic branching, neurogenesis, and synaptic density (for review, see (Ohline & Abraham, 2019; Van Praag et al., 2000). Probing different dimensions of experience common to poverty (e.g., stress exposure; environmental stimulation) to understand the patterns reported here will be critical moving forward (Palacios-Barrios et al., 2021; Palacios-Barrios & Hanson, 2019). In our supplement, we examined income and parental education independently and these are important sets of results to also consider. Thoroughly isolating major drivers of neurobiological and behavioral differences could be particularly powerful, especially if this information can be translated into effective interventions to lessen different SES-related disparities.

These limitations notwithstanding, our results provide data about neurobiological alterations seen in relation to SES. Few investigations have examined subdivisions of the amygdala and hippocampus in relation to SES, and our study represents the first to examine these associations with statistical interactions testing age and sex moderation in a large pediatric sample. Such neurobiological differences may connect to SES-gradients in health and well-being. These patterns also suggest how structural inequalities, in education, healthcare access, housing stability, and economic opportunity, may shape brain development, highlighting the critical need for policy interventions that address these systemic barriers. Additional research is needed to clarify the complex relations among early poverty exposure and long-term mental health difficulties; our data are, however, a needed step in the ability to understand the impact of SES on neurobiology critical for emotion, memory, and learning.

## Supporting information

Supplemental Materials/Analyses

## Author contributions

Jamie Hanson conceived of the project, outlined approaches to statistical analyses, and wrote the majority of the manuscript. Dorthea Adkins wrote portions of the manuscript and constructed statistical models. Brendon Nacewicz guided the team on conceptual and methodological elements of amygdala neurobiology and provided feedback on draft writing. Kelly Barry aided with data cleaning, coding, and organization.

