## Supplemental Materials/Analyses for "Socioeconomic Status and Subdivisions of the Amygdala and Hippocampus in Children and Adolescents"

**Supplemental Materials for “*****Influence of Socioeconomic Status on Amygdala and Hippocampus Subdivisions in Children and Adolescents*”**

I. *Additional Information about Participants in the Sample*

Participants between 5 and 18 years of age were drawn from four large, open-source neuroimaging projects: The National Consortium on Alcohol and Neuro-Development in Adolescence (NCANDA; (Brown et al., 2015), the Healthy Brain Network (HBN; (Alexander et al., 2017)), the Pediatric Imaging, Neurocognition, and Genetics (PING; (Jernigan et al., 2016), and the Human Connectome Project in Development (HCP-D; (Somerville et al., 2018). NCANDA recruited participants in a study examining developmental and biopsychosocial factors that predict susceptibility to substance abuse and the effects which substance use has on psychological and neurological development (Brown et al., 2015). HBN is an ongoing project focused on producing a large-scale study of healthy neurological and psychological development (Alexander et al., 2017). Recruitment for this project is still on-going, our data is drawn from waves 1 through 9. PING focused on exploring the relationships between neurological, genetic, and psychological factors impacting developmental trajectories (Jernigan et al., 2016). Only a portion of the original MRI data (i.e., non-derived values) were shared with research groups approved for public data use. HCP-D is an on-going study focused on understanding socioemotional, neurobiological, and environmental influences on mental health during sensitive developmental periods (Somerville et al., 2018). We were able to access a subset of participant data that included structural MRI volumes and demographic variables of interest. Across these combined studies, the total number of participants with demographic data and usable structural MRI volumes was N=2765 (44% Female; Mean Age=11.9, Age SD=3.5; Age Range=5.04-17.99).

II. *Information about Measurement of Socioeconomic Status*

Family or household income was assessed across all four studies, though question wording, reference period, and response format varied. Additional details about each study's income measure, including exact question wording and response options, are provided in Table S1. Notably, no study explicitly named different government benefit programs (e.g., Temporary Assistance for Needy Families; Supplemental Nutrition Assistance Program) as an includable income source. Given these measurement differences, income values were standardized within each study prior to analysis.

*— Insert Table S1 Here —*

III. *Detailed Information About MRI Acquisition*

High-resolution T1-weighted structural images were acquired with varying parameters across each project. For NCANDA, data was acquired across 5 sites: Duke University, University of Pittsburgh Medical Center (UPMC), Oregon Health & Science University (OHSU), University of California, San Diego (UCSD), and SRI International (SRI). All scans came from 3T scanners with 1.2x0.9375x0.9375mm resolution with a mixture of scanner manufacturers across Philips, Siemens, and General Electric. For HBN, data was acquired across 4 sites: Rutgers, Staten Island, CitiGroup Cornell Brain Imaging Institute, and City University of New York. The scans from Rutgers were collected on a Siemens 3 Tesla with a 0.8x0.8x0.8mm resolution. The scans from Staten Island were collected on a Siemens 1.5 Tesla with a 1.0x1.0x1.0mm resolution. The scans from Cornell were collected on a Siemens 3T scanner with a 1.0x1.0x1.0mm resolution. The scans from the City University of New York were collected on a Siemens 3 Tesla with a 0.8x0.8x0.8mm resolution. For PING, data was acquired across 8 sites. All scans came from 3T scanners with 1.0x1.0x1.0mm resolution with a mixture of scanner manufacturers across Philips, Siemens, and General Electric. For HCPD, data was acquired across 4 sites: Harvard University, University of California-Los Angeles, University of Minnesota, and Washington University in St. Louis. Across study sites, all scans came from Siemens 3T with a 0.8x0.8x0.8mm resolution for T1w images.

IV. *Information about MR Imaging Processing (Freesurfer).*

Standard-processing approaches from Freesurfer (e.g., cortical reconstruction; volumetric segmentation) were performed in version 7.1. Freesurfer is a widely documented and freely available morphometric processing tool suite (<http://surfer.nmr.mgh.harvard.edu/>) and was implemented using Brainlife.io, (brainlife.app.0, <https://doi.org/10.25663/bl.app.0>), which is a free, publicly funded, cloud-computing platform for reproducible neuroimaging pipelines and data sharing (Hayashi et al., 2024), for additional information, visit <http://brainlife.io/>). The technical details of this method are described in prior publications (Dale et al., 1999; Fischl et al., 2002, 2004; Fischl & Dale, 2000). Briefly, this processing includes motion correction and intensity normalization of T1-weighted images, removal of non-brain tissue using a hybrid watershed/surface deformation procedure, automated Talairach transformation, segmentation of the subcortical white matter and deep gray matter volumetric structures (including hippocampus, amygdala, caudate, putamen, ventricles), tessellation of the gray matter white matter boundary, and derivation of cortical surface area and cortical thickness. Of note, the "recon-all" pipeline with the default set of parameters (no flag options) was used and no manual editing was conducted. In keeping with our past work (Gilmore et al., 2021), Freesurfer outputs were checked via research staff for major errors, and via automated methods (see “*Image Quality Metrics*” in the main manuscript). After successful processing and quality assurance, we extracted volumes for our subcortical structures of interest– the hippocampus and amygdala. FreeSurfer version 7.1 natively includes options to segment both structures. Previous work has shown that the use of FreeSurfer 7 to segment hippocampal and amygdalar subregions yields at least good, if not excellent numerical reliability in all subregions (Kahhale et al., 2023). This method also yields at least good spatial reliability in all but one (left hippocampal fissure) hippocampal subfields and most amygdalar subnuclei except for the Medial and Paralaminar Nuclei.

The hippocampal segmentation method (Iglesias et al., 2015) is based on a hippocampal atlas initially produced from a dataset of 15 hand-traced high definition ex-vivo T1-weighted 7 T scans. The atlas for this method partitions the hippocampus into the following 12 subfields: (1) Parasubiculum, (2) Presubiculum [Head and Body], (3) Subiculum [Head and Body], (4) CA1 [Head and Body], (5) CA3 [Head and Body], (6) CA4 [Head and Body], (7) Granule Cell and Molecular Layer of the Dentate Gyrus [GC-ML-DG, Head and Body], (8) Molecular layer [Head and Body], (9) Fimbria, (10) Hippocampal Fissure, (11) Hippocampal Tail, and (12) Hippocampus-Amygdala-Transition-Area (HATA).

For the amygdala, the automated segmentation method is based on an atlas produced from 10 hand-traced high definition ex-vivo T1w 7T scans (Saygin et al., 2017). The amygdala atlas partitions the structure into the following 6 subnuclei: (1) Lateral, (2) Basal, (3) Central, (4) Medial, (5) Cortical, (6) Accessory Basal, (6) Paralaminar. Two additional subdivisions, the Corticoamygdaloid Transition Area and Anterior Amygdaloid Area, are also output. For both the hippocampal and amygdalar atlases, manually segmented ex-vivo data was applied to the probabilistic classification of the nodes on a parameterized deformation mesh of each respective region. Both hippocampal and amygdalar images were classified using a parameterized generative model and by optimizing the matching of voxels to its respective region in a Bayesian inference framework (for additional information, see (Iglesias et al., 2015).

V. *Notes regarding Image Quality*

Past work from our group (e.g., (Gilmore et al., 2021) has found that T1-weighted image quality is related to volumetric measures from commonly used morphometric tools suites (e.g., Freesurfer). We therefore assessed image quality to: 1) Exclude particularly high-motion scans; and 2) limit the impact of image quality on subcortical volume quantification. To assess MRI quality, we generated a quantitative metric (“CAT12 score”) using the Computational Anatomy Toolbox 12 (CAT12; (Gaser et al., 2024). This metric considers four summary measures of image quality: noise-to-contrast ratio, coefficient of joint variation, inhomogeneity-to-contrast ratio, and root-mean-squared voxel resolution. CAT12 normalizes and combines these measures using a kappa statistic-based framework. The score is a value from 0 to 1, with higher values indicating better image quality. Additional information is available at: <http://www.neuro.uni-jena.de/cat/index.html#QA>.

VI. *Measures of Socioeconomic Status (SES).*

While there was variability in how each study measured SES, we aimed to have a multimethod determination of this construct; specifically, we sought to use metrics of both parental education and household income. For parental education, each project asked parents and caregivers to report on how much school they completed (e.g., obtained a high school diploma; some college; graduate degree). We converted these self-reports into numbers of years (i.e., high school diploma=12 years; some college=14 years) and then took the highest level of education achieved by either parent or caregiver. For household income, parents or caregivers selected an income range (i.e., $30,000-39,999/year; $50,000-$99,999/year) or reported a continuous value. With income ranges and in keeping with best, past practices (Hanson et al., 2011), we took the midpoint of the range and then log-transformed this value ($30,000-39,999=$34,999.5; log($34,999.5)=4.54406184). For continuous income reports, we also log-transformed these values. Then, we created a SES composite by combining log-transformed income and maximum parental education. For this composite, each variable was first z-scored across all participants and z-scores from log-transformed income and maximum parental education were then averaged. Across all studies, income and education were found to be moderately correlated (r=0.45, p <.005).

VII. *Relations between Income and Volumes in Amygdala and Hippocampal Subdivisions*

We constructed linear mixed effects models to examine whether household income alone was related to hippocampal or amygdalar volumes. For each model, standardized hippocampal or amygdala subdivision volumes were entered as the dependent variable. Income-to-needs ratio was entered as an independent variable (fixed effect), along with age, sex (binary-coded), age^2^, image quality, and estimated total intracranial volume, while site was entered as a random effect.

Similar to our SES composite, income was positively associated with volumes across all three amygdala subdivisions. For the amygdala, higher income was related to larger volumes in the superficial cortical division (β=0.20, p<0.001, p-fdr<0.001), basolateral complex (β=0.17, p<0.001, p-fdr=0.002), and centromedial region (β=0.12, p=0.028, p-fdr=0.042). Among hippocampal subfields, income was significantly related to volumes in the head of the hippocampus (β=0.17, p=0.001, p-fdr=0.002), with no significant associations observed for the body (p=0.656, p-fdr=0.656) or tail (p=0.422, p-fdr=0.507) after correcting for multiple comparisons. No significant Income x Sex, Income x Age, or Income x Sex x Age interactions emerged for any subdivision after FDR correction (all p-fdr>0.58).

*— Insert Figure S1 Here —*

VIII. *Associations between Parental Education and Volumes in Amygdala and Hippocampal Subdivisions*

We constructed linear mixed effects models to examine whether maximum parental education alone was related to hippocampal or amygdalar volumes. For each model, standardized hippocampal or amygdala subdivision volumes were entered as the dependent variable. Maximum parental education (in years) was entered as an independent variable (fixed effect), along with age, sex (binary-coded), age^2^, image quality, and estimated total intracranial volume, while site was entered as a random effect.

Parental education was positively associated with volumes in two of the six subdivisions examined. Higher parental education was related to larger volumes in the amygdala superficial cortical division (β=0.023, p=0.002, p-fdr=0.014) and the hippocampal head (β=0.022, p=0.005, p-fdr=0.014). While uncorrected analyses suggested parental education was also associated with the amygdala basolateral complex (β=0.015, p=0.033) and centromedial region (β=0.016, p=0.043), these effects did not survive FDR correction (p-fdr=0.064 for both). No significant associations were observed for the hippocampal body (p=0.624, p-fdr=0.683) or tail (p=0.683, p-fdr=0.683).

No significant Parental Education x Sex, Parental Education x Age, or Parental Education x Sex x Age interactions emerged for any subdivision after FDR correction (all p-fdr>0.23). These results suggest that the association between parental education and subcortical volumes was consistent across sex and age in this sample.

*— Insert Figure S2 Here —*

IX. *Associations with SES and Volumes in Amygdala and Hippocampal Subdivisions (Using Canonical Subnuclei and Subfields Output by Freesurfer 7)*

Linear mixed effects models were run to examine the effect of SES on the canonical amygdala subnuclei and hippocampal subfields output by Freesurfer. An SES composite score was computed by first z-scoring both the income-to-needs ratio and maximum parental education variables, and then averaging them together for each participant. For these analyses, we created linear mixed effects models with standardized regional volumes as dependent variables and the SES composite (along with age, sex, age², image quality, and estimated total intracranial volume) as independent variables (fixed effects), while site was entered as a random effect.

Higher SES was positively associated with volumes in six of the nine amygdala subnuclei examined. SES showed the strongest associations with the anterior amygdaloid area (β=0.10, p<0.001, p-fdr=0.001), accessory basal nucleus (β=0.075, p<0.001, p-fdr=0.004), and basal nucleus (β=0.075, p<0.001, p-fdr=0.004). SES was also positively related to volumes in the corticoamygdaloid transition area (β=0.063, p=0.005, p-fdr=0.015), central nucleus (β=0.062, p=0.008, p-fdr=0.022), and paralaminar nucleus (β=0.058, p=0.009, p-fdr=0.022). SES was not significantly related to volumes in the cortical nucleus, medial nucleus, or lateral nucleus after correcting for multiple comparisons.

In the hippocampus, higher SES was associated with larger volumes in multiple subregions within the hippocampal head. Specifically, SES was positively related to presubiculum head (β=0.087, p<0.001, p-fdr=0.004), subiculum head (β=0.079, p=0.002, p-fdr=0.011), CA1 head (β=0.065, p=0.005, p-fdr=0.015), and molecular layer head (β=0.064, p=0.005, p-fdr=0.015) volumes. SES was also positively associated with the hippocampus-amygdala transition area (HATA; β=0.076, p=0.002, p-fdr=0.009), fimbria (β=0.059, p=0.015, p-fdr=0.034), and parasubiculum (β=0.062, p=0.020, p-fdr=0.039).

Notably, SES showed negative associations with two hippocampal body subregions: CA4 body (β=-0.056, p=0.020, p-fdr=0.039) and CA3 body (β=-0.059, p=0.025, p-fdr=0.046), such that higher SES was related to smaller volumes in these regions. SES was not significantly related to volumes in the granule cell and molecular layer of the dentate gyrus (GC-ML-DG), CA3 head, CA4 head, CA1 body, subiculum body, presubiculum body, molecular layer body, or hippocampal fissure after FDR correction.

No significant SES x Sex, SES x Age, or SES x Sex × Age interactions emerged for any amygdala or hippocampal subdivision after FDR correction (all p-fdr>0.73). These results suggest that the associations between SES and Freesurfer-defined subdivisions were consistent across sex and age in this sample.

*— Insert Figure S3 Here —*

X. ***Analysis Probing Non-Linear Age Effects Using Generalized Additive Mixed Models***

In our original models, we examined interactions between age, sex, and SES, and used age^2^ as a covariate. But, age could have non-linear interactive effects, and this may require additional statistical models. It is, however, unclear whether one should include interactions with age (with a quadratic effect) and age^2^ in interactions in these types of models. We therefore conducted a sensitivity analysis using generalized additive mixed models (GAMMs; mgcv; (Wood, 2025), replacing the age terms with a regression spline (k=5) and testing whether this non-linear age trajectory was moderated by sex and SES via varying coefficient smooths, with site as a random effect. The three-way age-by-SES-by-sex interaction and all lower-order age interaction terms were tested using approximate F-tests with FDR correction applied across regions. SES main effects were extracted as parametric estimates in models where SES was included as a standard linear term alongside the age spline. In these models, SES main effects replicated the primary findings: higher SES was associated with larger volumes in all three amygdala subdivisions (basolateral: β=0.042, q=.006; centromedial: β=0.048, q=.006; superficial: β=0.055, q < .001) and the hippocampal head (β=0.076, q < .001), with no significant effects for hippocampal body or tail. These are shown in Figure S4. The three-way age-by-SES-by-sex interaction smooth was non-significant in all six regions (all F < 1.76, all q > .60), and the two-way age-by-SES interaction was non-significant after FDR correction in all regions except the hippocampal head (F=8.47, q=.001). These results are broadly consistent with the primary analyses noted in the main manuscript, especially the absence of significant three-way interactions.

*— Insert Figure S4 Here —*

XI. ***Cross-Validation of Statistical Models from Main Document***

To test whether SES effects generalize across acquisition sites, we conducted leave-one-site-out cross-validation (LOSO-CV) across 21 sites (total N=2,765). Each fold withheld one site, estimated model parameters on the remaining data, and generated fixed-effects-only predictions for held-out observations. Because site conflates scanner hardware, acquisition protocol, and recruitment context, this is a good test of replicability.

We fit four nested models at each fold: M0 (covariates only: age, age², TIV, scan quality), M1 (+ SES and sex main effects), M2 (+ all two-way interactions), and M3 (+ three-way SES × sex × age). All volumes were standardized, so coefficients are in SD units.

At each fold we extracted the fixed-effect SES estimate, yielding a distribution of 21 betas, one per site. We tested whether this distribution differed from zero via one-sample t-tests, FDR-corrected across regions. A nonzero mean with low fold-to-fold variance indicates the SES effect is consistent regardless of which site is withheld - which the in-sample p-value cannot establish. We also computed fold-level ΔR²=R²(M1) - R²(M0) and tested its mean against zero; a consistently positive out-of-sample ΔR² rules out overfitting as an explanation. As a site-level check, we correlated SES with M0 residuals within each held-out site. All analyses used R 4.5.2 (R Core Team, 2025), lme4 2.0.1 (Bates et al., 2015), and lmerTest 3.2.1 (Kuznetsova et al., 2017).

Across 21 LOSO-CV folds, SES was positively associated with volume in all three amygdala subdivisions and hippocampal head in every fold (100% of folds, all FDR q<.001). Mean fold-level betas ranged from 0.042 SD (basolateral amygdala) to 0.076 SD (hippocampal head) per SD increase in SES, with cross-fold SDs of 0.003-0.006. Hippocampal body showed no reliable association (mean β=0.001, q=.105). Hippocampal tail was statistically significant but directionally unstable: the mean beta was negative (β=−0.004, q < .001), yet only 10% of folds produced a negative coefficient, making the result difficult to interpret. These SES effects are shown in Figure S5.

*— Insert Figure S5 Here —*

**Table S1. Comparison of Family/Household Income Questions Across Developmental Neuroimaging Datasets**

| **Study** | **Variable name** | **Exact question wording** | **Reporter** | **Reference period** | **Format** | **Brackets / range** |
| --- | --- | --- | --- | --- | --- | --- |
| **NCANDA**  Brown et al., 2015 | parentreport_sesp14 | *"Which of these categories best describes your* ***TOTAL COMBINED FAMILY INCOME*** *for the past 12 months? This should include income (before taxes and deductions) from all sources, wages, rent from properties, social security, disability and/or veteran’s benefits, unemployment benefits, workman’s compensation, help from relatives (including child payments and alimony), and so on."* | Parent/guardian | Past 12 months | Dropdown; 11 options | <$5K \| $5K–11,999 \| $12K–15,999 \| $16K–24,999 \| $25K–34,999 \| $35K–49,999 \| $50K–74,999 \| $75K–99,999 \| $100K–199,999 \| $200K+ \| Don’t know |
| **HBN**  Alexander et al., 2017 | FSQ_04 | *"What is your annual household income?"* | Parent/guardian | Annual | Ordinal; 13 options | <$10K \| $10–19,999 \| $20–29,999 \| $30–39,999 \| $40–49,999 \| $50–59,999 \| $60–69,999 \| $70–79,999 \| $80–89,999 \| $90–99,999 \| $100–149,999 \| $150K+ \| Choose not to disclose |
| **PING**  Jernigan et al., 2016 | FDH_3_Household_Income | Full question wording not available, but “*Parents were also asked to report the total yearly family income*.” (per Noble et al., 2015) | Parent/guardian or self-report (≥18 yrs) | Total yearly income | Ordinal; 12 categories | <$5K \| $5–9,999 \| $10–19,999 \| $20–29,999 \| $30–39,999 \| $40–49,999 \| $50–99,999* \| $100–149,999 \| $150–199,999 \| $200–249,999 \| $250–299,999 \| $300K+ |
| **HCP-D**  Somerville et al., 2018 | annual_fam_inc (socdem01) | *"The following questions are about your family and household. Please state your* ***TOTAL COMBINED FAMILY INCOME*** *for the past 12 months. This should include income (before taxes and deductions) from all sources, wages, rent from properties, social security, disability and/or veteran’s benefits, unemployment benefits, workman’s compensation, help from relatives (including child payments and alimony), and so on."* | Caregiver | Past 12 months | Continuous (open dollar entry) | No brackets. Raw dollar value entered by respondent. Researcher applies any binning post-hoc. |

Figure S1.

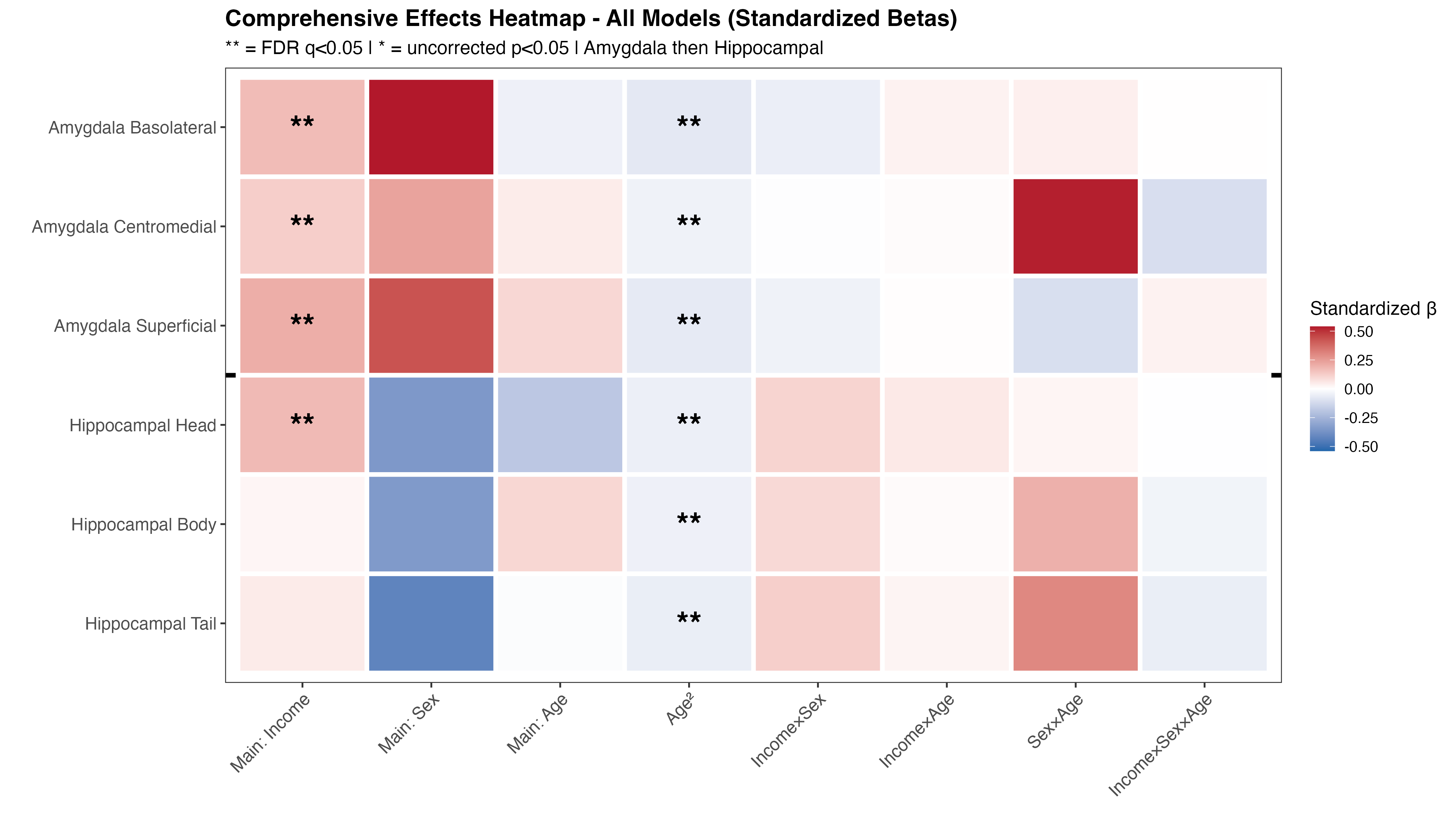

Caption: Standardized β coefficients from linear mixed-effects models predicting amygdala subdivision (basolateral complex, centromedial, superficial cortical) and hippocampal subfield (head, body, tail) volumes. Predictors included income, sex, age, age², and two- and three-way interactions; models controlled for image quality and intracranial volume, with site as a random effect. Red = positive association, blue = negative; ** FDR q < 0.05, * uncorrected p < 0.05.

Figure S2.

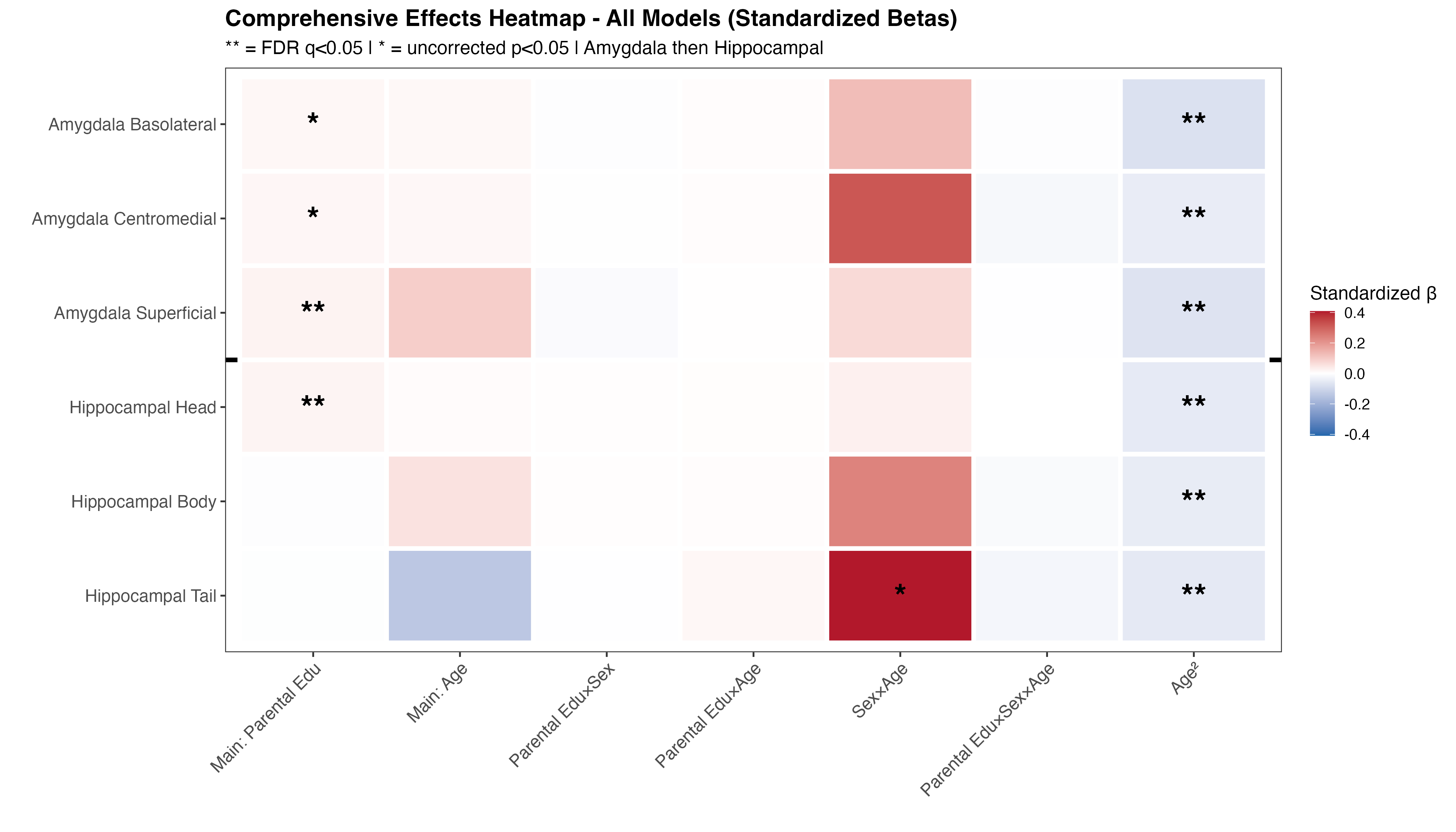

Caption: Standardized β coefficients from linear mixed-effects models predicting amygdala subdivision (basolateral complex, centromedial, superficial cortical) and hippocampal subfield (head, body, tail) volumes. Predictors included caregiver education, sex, age, age², and two- and three-way interactions; models controlled for image quality and intracranial volume, with site as a random effect. Red = positive association, blue = negative; ** FDR q < 0.05, * uncorrected p < 0.05.

Figure S3.

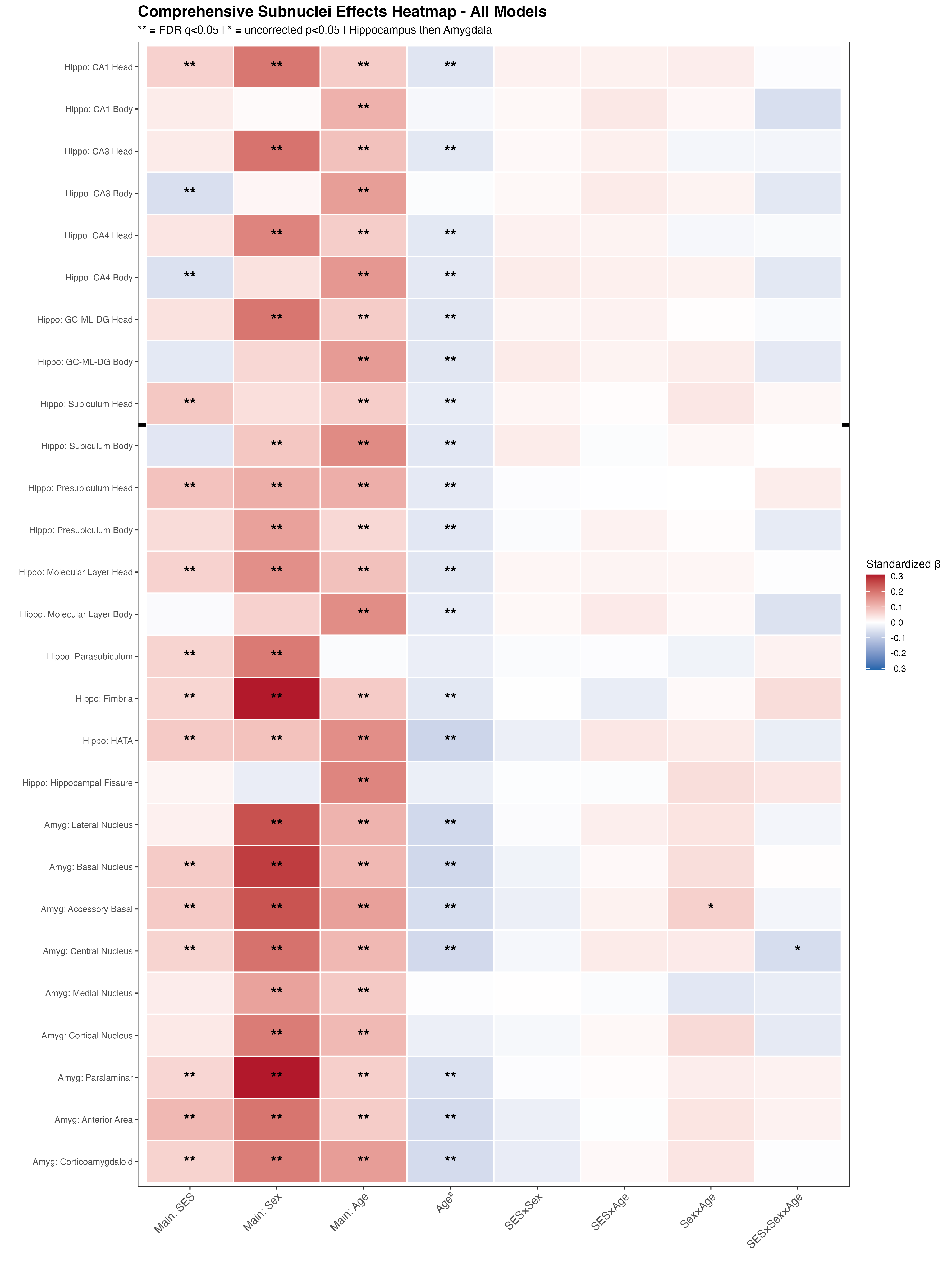

Caption: Standardized β coefficients from linear mixed-effects models predicting volumes across 9 amygdala subnuclei and 18 hippocampal subfields as defined by FreeSurfer 7. The SES composite was the mean of z-scored income-to-needs ratio and maximum parental education. Models controlled for age, sex, age², image quality, and intracranial volume, with site as a random effect. Red = positive association, blue = negative; ** FDR q < 0.05, * uncorrected p < 0.05.

Figure S4.

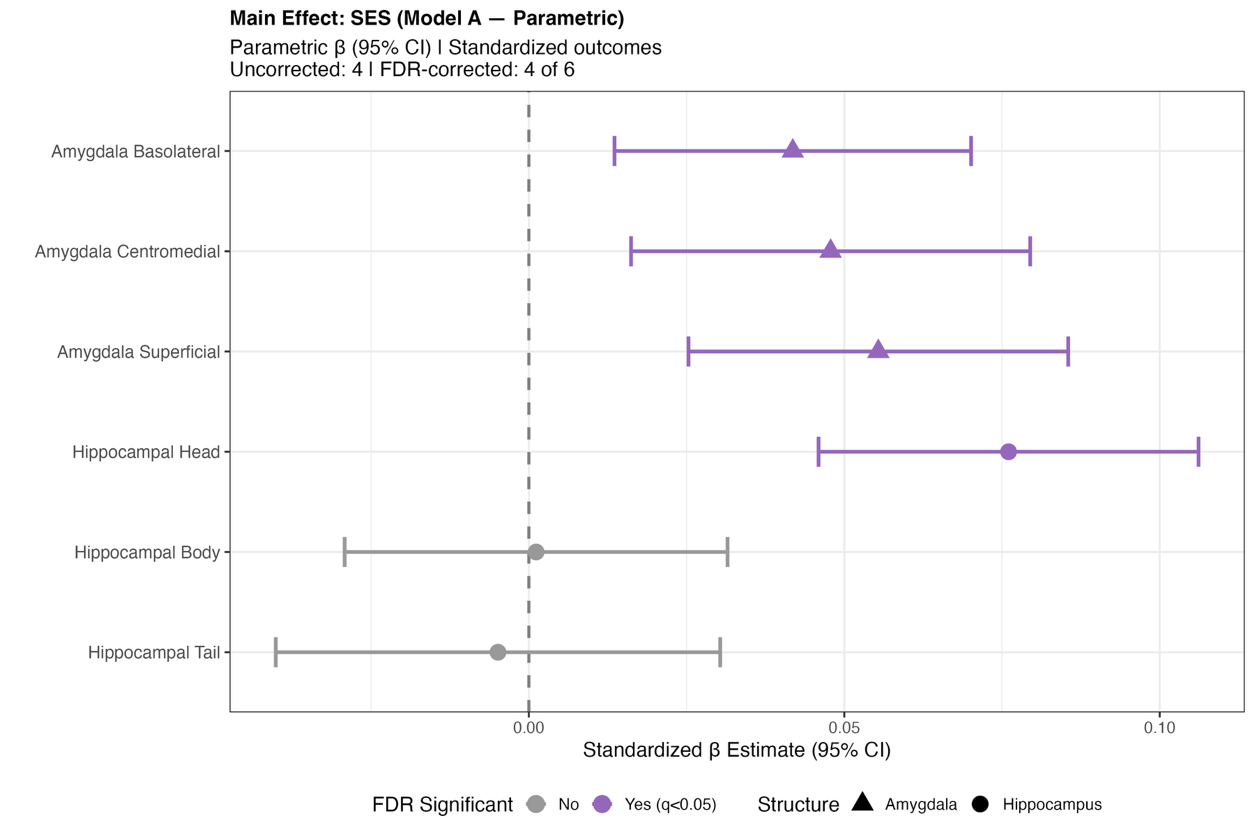

Caption: Parametric β estimates (95% CI) for the SES composite from GAMMs replacing the quadratic age term with a regression spline (k = 5), with site as a random effect. Purple = FDR-significant (q < 0.05); gray = non-significant. Triangles = amygdala subdivisions; circles = hippocampal subfields.

Figure S5.

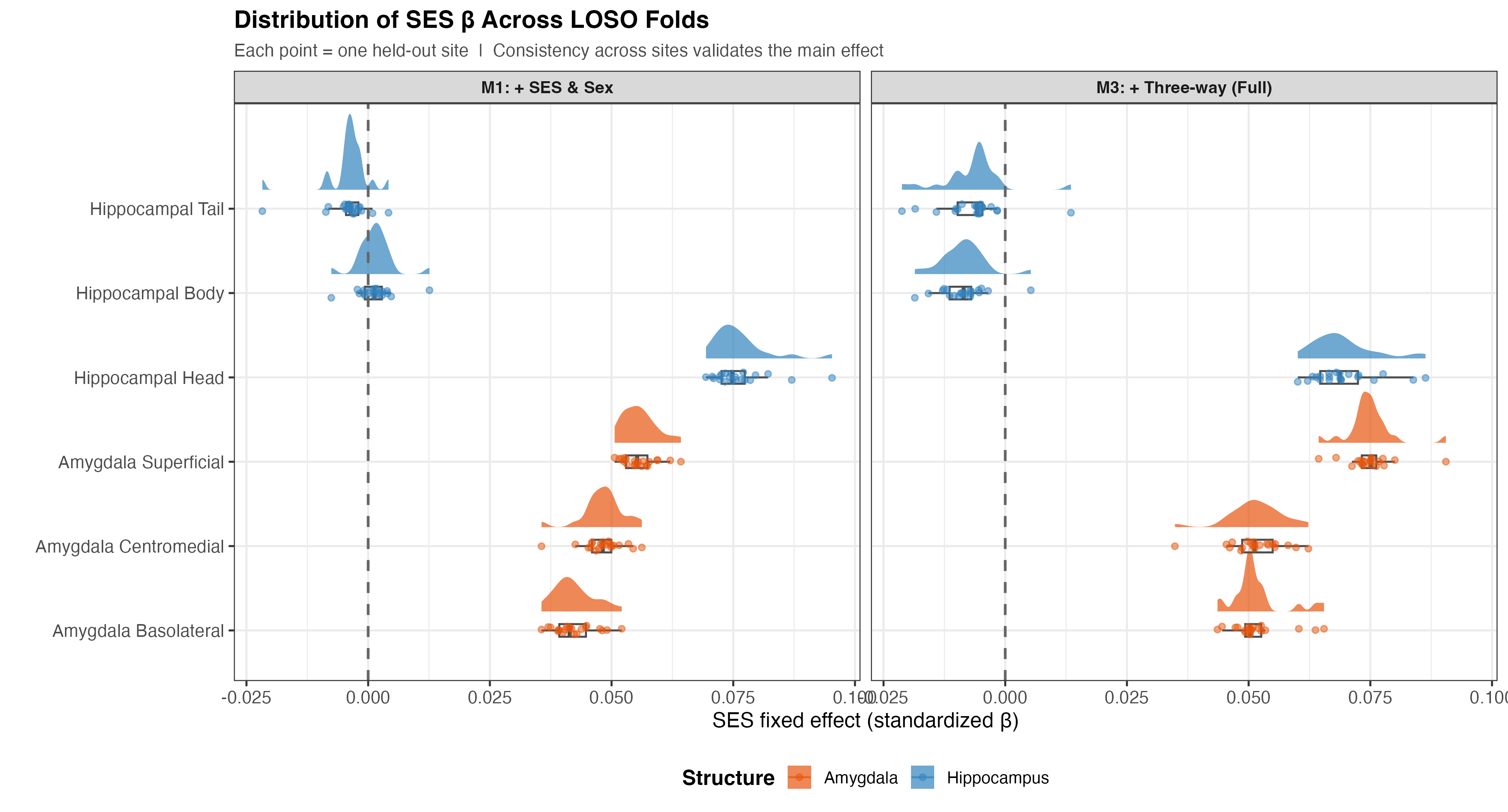

Caption: Distribution of fixed-effect SES β estimates (standardized) across 21 LOSO-CV folds for Model 1 (SES and sex main effects) and Model 3 (full three-way interaction). Each point is one held-out site. Orange = amygdala subdivisions; blue = hippocampal subfields. Dashed line at zero.
